# TPR Domains Drive the Functional Phase Separation of HOP and its Regulation by Hsp90 and Hsp70

**DOI:** 10.64898/2026.06.05.730378

**Authors:** Benjamin S. Rutledge, Bryce E. Gnidec, Marko Vukcevic, Solomon Hussein, Desmond Prah Amoah, Paul LaPointe, Marco A.M. Prado, Wing-Yiu Choy, Martin L. Duennwald

## Abstract

HOP is a cochaperone that facilitates client transfer between two major chaperones, Hsp90 and Hsp70. Emerging evidence, however, suggests that HOP plays additional roles in coordinating complex proteostasis networks. Upon exposure to proteostatic stress, HOP rapidly sequesters soluble misfolded proteins into cytoplasmic foci in a Hsp90 independent manner, thereby facilitating their clearance through the ubiquitin proteasome system. We demonstrate here that stress-dependent HOP foci are biomolecular condensates formed by liquid-liquid phase separation. Purified HOP forms protein droplets that closely resemble the foci observed in cells. Our biophysical analyses show that the phase separation of HOP is driven by electrostatic interactions between its tandem TPR domains, with a critical role of its TPR2A domain. Of note, Hsp90 and Hsp70 regulate the extent of HOP phase separation, with Hsp70 driving HOP droplet formation and Hsp90 reversing it. Finally, we find that the Y354E phosphomimetic variant of HOP impairs phase separation and sensitizes cells to acute misfolding stress, suggesting a key role of HOP condensation in mitigating protein misfolding stress. Our work thus identifies a new mechanism by which HOP phase separation mitigates protein misfolding stress in eukaryotic cells and is regulated by Hsp70 and Hsp90.

## Introduction

Hsp-organizing protein (HOP, also known as Stress inducible phosphoprotein 1, STIP1 and Sti1 in yeast) is a major co-chaperone of both Hsp90 and Hsp70, as it is one of the few proteins that directly interact with both chaperones and facilitate the formation of the Hsp90-HOP-Hsp70 complex[1]. In addition, HOP interacts with many other proteins involved in proteostasis[2], acting as a hub to coordinate protein folding, refolding, and degradation[3]. The deletion of HOP in higher order organisms, such as mice, is lethal[4]. However, yeast cells deleted for Sti1, the yeast homolog of HOP, exhibit no negative growth phenotype under optimal growth conditions[5]. We demonstrated that during proteostatic stress, Sti1 and HOP form cytoplasmic foci in yeast and mammalian cells, respectively. These cytoplasmic foci sequester misfolded proteins[3], and the deletion of STI1 prevents the formation Hsp90/Hsp70/Hsp104 and proteasome-dependent heat-induced inclusions[6]. Once stress subsides, cytoplasmic Sti1 foci are rapidly cleared, a process that relies on the foldase activity of Hsp90[3].

Here, we demonstrate that cytoplasmic foci formed by Sti1 or HOP are biomolecular condensates. Biomolecular condensates are membraneless assemblies containing high local concentrations of specific proteins and other biomolecules formed through liquid-liquid phase separation[7]. The subcellular localization and macromolecular crowding within condensates can accelerate specific enzymatic reactions and modulate other biological functions[8,9]. For instance, the recruitment of proteostasis components to specialized condensates is required for the proper function of the ubiquitin-proteasome system[10]. Interestingly, recent machine learning models predict that HOP, alongside other proteostasis proteins such as Hsp90, UBQLNs, RAD23, and proteasome subunits, form biomolecular condensates[11]. Notably, only a small fraction of Sti1 foci co-localize with other established biomolecular condensates such as stress granules, suggesting that Sti1 condensates are distinct and can form independently of these structures[3,12]. Yet, overall, the molecular and cellular mechanisms driving Sti1 foci formation under proteostasis stress remain largely unknown.

Like other biomolecular condensates that form under stimulating or stress conditions, Sti1/HOP foci spontaneously assemble in response to protein misfolding stress and are reversibly dissolved when the stress subsides[3,13]. Even though the ability to phase separate is commonly attributed to intrinsically disordered regions (IDRs) of low sequence complexity[14], folded domains emerge as central regulators of phase separation[15]. HOP is composed of two low complexity DP domains, and three structured tetratricopeptide repeat (TPR) domains[16]. The three TPR domains (TPR1, TPR2A, and TPR2B) of HOP mediate its interactions with Hsp90 and Hsp70, with TPR2A binding Hsp90 and the TPR1 and TPR2B domains interacting with Hsp70[1,16–18].

Notably, recent research using engineered TPR proteins indicates that the structural attributes and elongated nature of tandem TPR domains facilitate phase separation. Furthermore, short binding motifs within these TPR repeats enable the recruitment of clients into condensates formed by TPR proteins[19]. The tandem TPR2A and TPR2B domains in HOP adopt an elongated structure, sharing many structural characteristics with the engineered TPR proteins used to study phase separation[19,20].

Here, we show that Sti1/HOP phase separates into liquid droplets both *in vivo* and *in vitro.* Phase separation of HOP is driven by its TPR domains, mainly through electrostatic interactions. Furthermore, we demonstrate that the binding of Hsp90 to the TPR2A domain disassembles HOP droplets, while Hsp70 binding to TPR1 and TPR2B promotes the formation of larger droplets *in vitro* and *in vivo*. The phase separation of HOP can be further regulated by the presence of ATP, which binds both TPR2A and TPR2B. The phosphomimetic Y354E variant of HOP impairs its ability to undergo phase separation, sensitizing cells to acute misfolding stress. Our study suggests a previously unrecognized mechanism by which Hsp70- and Hsp90-dependent HOP-mediated phase separation facilitates the sequestration of misfolded proteins and their rapid refolding or degradation, which is key for cellular survival.

## Methods

### Yeast Strains

Yeast strains BY 4741 (MAT a his3Δ1 leu2Δ0 lys2Δ0 ura3Δ0) and W303 (MAT a leu2-3112 trp1-1 can1-100 ura3-1 ade2-1 his3-11,15 [phi+]) were used in this study. The *Δsti1* W303 yeast strain was produced as previously described[3]. Yeast expressing the C-terminal GFP fusion protein of Sti1 (Sti1-GFP) under its endogenous promoter were obtained from the Schuldiner laboratory Swap-Tag method (SWAT) library[21].

### Yeast Media

Yeast-peptone-dextrose (YPD)-rich media (10 g/L of yeast extract, 20 g/L of peptone, and 20 g/L of dextrose) and selective dextrose (SD) media (6.7 g/L yeast nitrogen base (YNB), 60 mg/L of L-isoleucine, 20 mg/L of L-arginine, 40 mg/L of L-lysine HCl, 60 mg/L of L-phenylalanine, 10 mg/L of L-threonine, and 2% glucose) in either liquid media or agar plates (20 g/L) were used to grow yeast cells. SD media was supplemented with four different amino acids (40 mg/L of L-tryptophan, 60 mg/L of L-leucine, 20 mg/L of L-histidine-monohydrate, 10 mg/L of L-methionine, and 20 mg/L of uracil) depending on the selective marker of the plasmid. To expose cells to 1,6-Hexanediol, liquid media was supplemented with 1,6-Hexanediol to a final concentration of either 2.5% or 5%.

### Yeast Transformations

Yeast transformations were performed utilizing the standard PEG/lithium acetate transformation method[22].

### Fluorescence Recovery After Photobleaching

SD media was inoculated with yeast expressing fluorescently tagged constructs and incubated overnight at 30 °C in a shaking incubator. For each sample, 1 mL of culture was placed in an uncoated MATTEK 35 mm dish with a No. 1.5 coverslip that was pretreated with Concanavalin A to minimize cell movement. To induce heat-shock, the dishes were incubated at 42 °C for 1 hour prior to imaging unless otherwise indicated. Each culture was imaged using a Leica TCS SP8 microscope equipped with a 63 × 1.4 NA oil immersion objective. For these assays, a designated region of interest (ROI) was subjected to photobleaching for 3.6 seconds (11 frames at 0.328 seconds per frame) at 30% laser power, and the subsequent recovery of fluorescence intensity within the ROI was monitored and recorded for each experiment. The intensity recovery curves were then normalized and corrected for photobleaching effects.

### Recombinant Protein Expression and Purification

pDEST17 (Invitrogen) expression plasmids encoding full-length human HOP, TPR1 domain (TPR1; residues 1–118), TPR2A domain (TPR2A; residues 217–352), and TPR2B domain (TPR2B; residues 353–480) fused with N-terminal tobacco etch virus (TEV) cleavable hexa-His tags were transformed into the *E. coli* BL21 (DE3) pLysS strain for protein expression in minimal M9 media containing either ^14^NH_4_Cl (1 g/L) or ^15^NH_4_Cl (1 g/L) as the sole nitrogen source. Expression and purification of full-length human HOP and the TPR domains were performed as previously described[23]. pET151 bacterial expression vectors encoding human Hsp90 constructs were transformed into the *E. coli* BL21 strain. Expression and purification of human Hsp90 constructs were performed as previously described[24].

The pMCSG7 expression plasmid encoding full-length human Hsc70 (kindly gifted by Dr. Sue-Ann Mok, University of Alberta), fused with an N-terminal TEV cleavable hexa-His tag, was transformed into *E. Coli* BL21 (DE3) for expression in LB media. The expression and purification protocols for Hsc70 were adapted from those previously described[25]. In short, after overnight induction with 1 mM IPTG, cells were lysed and recombinant Hsc70 was purified by Nickel affinity chromatography using Ni Sepharose 6 Fast Flow beads (GE Healthcare Life Sciences). The 6xHis-tag was cleaved by incubation with 6xHis-tagged TEV overnight at 4 °C. The 6xHis-tagged TEV and cleaved 6xHis-tag were subsequently removed by additional Nickel affinity chromatography. The A_260_/A_280_ ratio of the purified protein was measured to confirm that the purified protein was free of nucleic acid contamination and purified Hsc70 was immediately frozen and stored at −80 °C. Protein concentrations were determined by measuring absorbance at 280 nm using a NanoDrop spectrophotometer.

### Fluorescence Imaging of Yeast Cells

SD media was inoculated with yeast expressing fluorescently tagged constructs and incubated overnight at 30 °C in a shaking incubator. To induce heat-shock, cultures were incubated at 42 °C for 1 hour unless otherwise indicated. Each culture was imaged using either a Zeiss Axio Vert.A1 microscope (1-2 μL placed on a microscope slide) or a Leica TCS SP8 microscope (1 mL placed in an uncoated MATTEK 35 mm dish with a No. 1.5 coverslip). For each condition, three biological replicates were imaged, and three images of random fields of each biological replicate were taken. Each data point represents the average percentage of cells that contained foci across the three images taken for each respective biological replicate. For each biological replicate, a minimum of 100 cells was counted towards the percentage of cells containing foci unless otherwise stated.

### Spectrophotometric Absorbance Measurements

Protein samples were prepared by mixing predetermined amounts of the proteins of interest and additives at desired concentrations in a base solution of 20 mM HEPES and 50 mM NaCl. The absorbance at 600 nm was monitored at ambient temperature using a Cytation 5 Cell Imaging Multi-Mode Reader (Biotek, Winooski, VT, USA) after the samples had incubated at ambient room temperature for 20 minutes after mixing. For time course measurements, the samples were monitored immediately after mixing for the indicated time intervals. When tracking the impact of temperature, the absorbance of each sample was measured 2 minutes after reaching temperature for every 2 °C increment. The absorbance values reported are the average of 3 biological replicates after subtracting the optical density of reference buffers.

### Fluorescent Labeling

Proteins were labelled with Alexa Fluor 680 (Alexa 680), Alexa Fluor 568 (Alexa 568), or Alexa Fluor 488 (Alexa 488) C5-maleimide dye (ThermoFisher Scientific). Dyes were used at a 1:1 molar ratio to their respective protein and mixtures were kept in dark conditions for a 30-minute incubation at room temperature. After incubation, the sample was buffer exchanged into fresh 20 mM HEPES and 50 mM NaCl at pH 7.2 using a 10 kDa MWCO Vivaspin Turbo 4 (Sartorius) concentrator with 4 sample volumes of fresh buffer to remove unreacted dye.

### Brightfield/Fluorescence Imaging of Phase Separation

Protein samples were prepared by mixing predetermined amounts of the proteins of interest and additives at desired concentrations in a base solution of 20 mM HEPES and 50 mM NaCl. Samples were either measured using the 20X objective lens on a Cytation 5 Cell Imaging Multi-Mode Reader (Biotek, Winooski, VT, USA) in black non-binding half-volume 96-well plates with clear bottoms (Corning Incorporated) or using the 63X objective lens on a Leica TCS SP8 microscope in an uncoated MATTEK 35 mm dish with a No. 1.5 coverslip. Unless otherwise indicated, samples were imaged 20 minutes after mixing and at ambient room temperature. To quantify droplet area and number, fluorescent microscopy of Alexa 680-labelled HOP was used to enhance the contrast between droplets and the background. Images were analyzed using the Analyze Particles function of ImageJ as 8-bit images with a circularity threshold of 0.7-1.0 and a minimum size threshold of 2 pixel^2^[26].

### Isolation of Phase-Separated Condensates from the Aqueous Phase

Protein condensates were isolated from the aqueous phase using an adapted protocol based on Xing et al.[27]. Protein samples were prepared by mixing predetermined amounts of the proteins of interest and additives at desired concentrations in a base solution of 20 mM HEPES and 50 mM NaCl. To remove potential contaminating condensates, protein stocks were centrifuged at 14,000 x *g* for 1 minute prior to use. Mixtures (40 μL final volume) were incubated for 20 minutes at room temperature, after which they were subject to centrifugation at 14,000 x *g* for 15 minutes at room temperature. The aqueous phase/supernatant (40 μL) was collected into a new tube, and the condensate/pellet was washed once with the sample’s corresponding buffer and resuspended in an equal volume as the supernate (40 μL). The supernate and pellet fractions were both redissolved with SDS-PAGE loading dye and loaded on a NuPAGE™ Bis-Tris Midi 4 to 12% Protein Gel (Invitrogen) and subsequently imaged on a ChemiDoc MP Imaging System (Bio-Rad) and analyzed using Image Lab (Bio-Rad).

### Solution NMR Spectroscopy

NMR experiments were performed on a 600 MHz Bruker spectrometer equipped with an xyz-gradient triple resonance probe. The ^1^H-^15^N HSQC spectra were collected using 128 × 537 complex points in the ^15^N and ^1^H dimensions, respectively. For experiments utilizing ^15^N labelled TPR domains, all experiments were conducted at 25 °C in 20 mM HEPES and 50 mM NaCl at pH 7.2. NMR data were processed using NMRFx Analyst[28] and analyzed using NMRViewJ[29] software. Chemical shift analyses of titration sets were conducted using the Titration Analysis function in NMRViewJ. Chemical shift perturbations for ^1^H-^15^N peaks were calculated following Δδ = [(Δδ_1H_)^2^ + (0.14*Δδ_15N_)^2^]^1/2^, where Δδ_1H_ and Δδ_15N_ are the changes in ^1^H and ^15^N chemical shifts (in ppm) upon the addition of the binding target[30].

### Growth Assays (Spotting Assays) and Quantification

The relative growth of the yeast cells was measured using spotting assays and quantification as previously described by Petropavlovskiy et al.[31]. In summary, 3 mL of SD media was inoculated with cells and incubated in a shaking incubator at 30 °C overnight. A 48-prong frogger (V&P Scientific, San Diego, CA, USA) was used to spot normalized serial dilutions of cells onto SD plates lacking the selective amino acid. When testing the response to acute heat shock, cell dilutions were incubated at 42 °C for one hour and immediately spotted onto plates. Plates were incubated at either 30 °C or 37 °C, and growth was monitored during the entire growth period. All spotting assays contain respective control lanes on each experimental plate, and images of the plates were cropped and separated to highlight relative side-by-side comparisons. Unless otherwise indicated, the pixel count of the third dilution of the growth assay was used for quantification. Values were normalized to the average of the respective control lanes. For all growth assays, a minimum of three biological replicates were used.

### Statistical analysis

Statistical analysis of condensation sedimentation assays, condensate droplet size/number, and percentage of foci assessed by microscopy were performed using the GraphPad Prism X software (GraphPad Software, Boston, MA). To determine statistical significance, unpaired one-way or two-way ANOVA tests were used to compare means and standard deviations between relevant controls and experimental data sets (each data set was composed of a minimum of 3 repeats). Error bars represent standard errors of the mean.

## Results

### Sti1 and HOP form liquid-like condensates in the cytoplasm during cells stress

We previously reported that protein misfolding stress causes Sti1 in yeast and HOP in human cells to localize into distinct cytoplasmic inclusions, which dissolve once the stress subsides[3]. The soluble and transient nature of Sti1/HOP foci led us to hypothesize that these localized clusters of Sti1/HOP are liquid droplets in nature. Supporting this idea, multiple sequence-based LLPS analysis tools[32–35] predict that Sti1 /HOP have a high propensity to undergo phase separation (Figure 1A). In contrast, most prediction tools predict that Aha1, another co-chaperone of Hsp90, does not phase separate (Figure 1A), consistent with our previous observations that these two co-chaperones display distinct foci formation behaviors[3].

**Figure 1.**
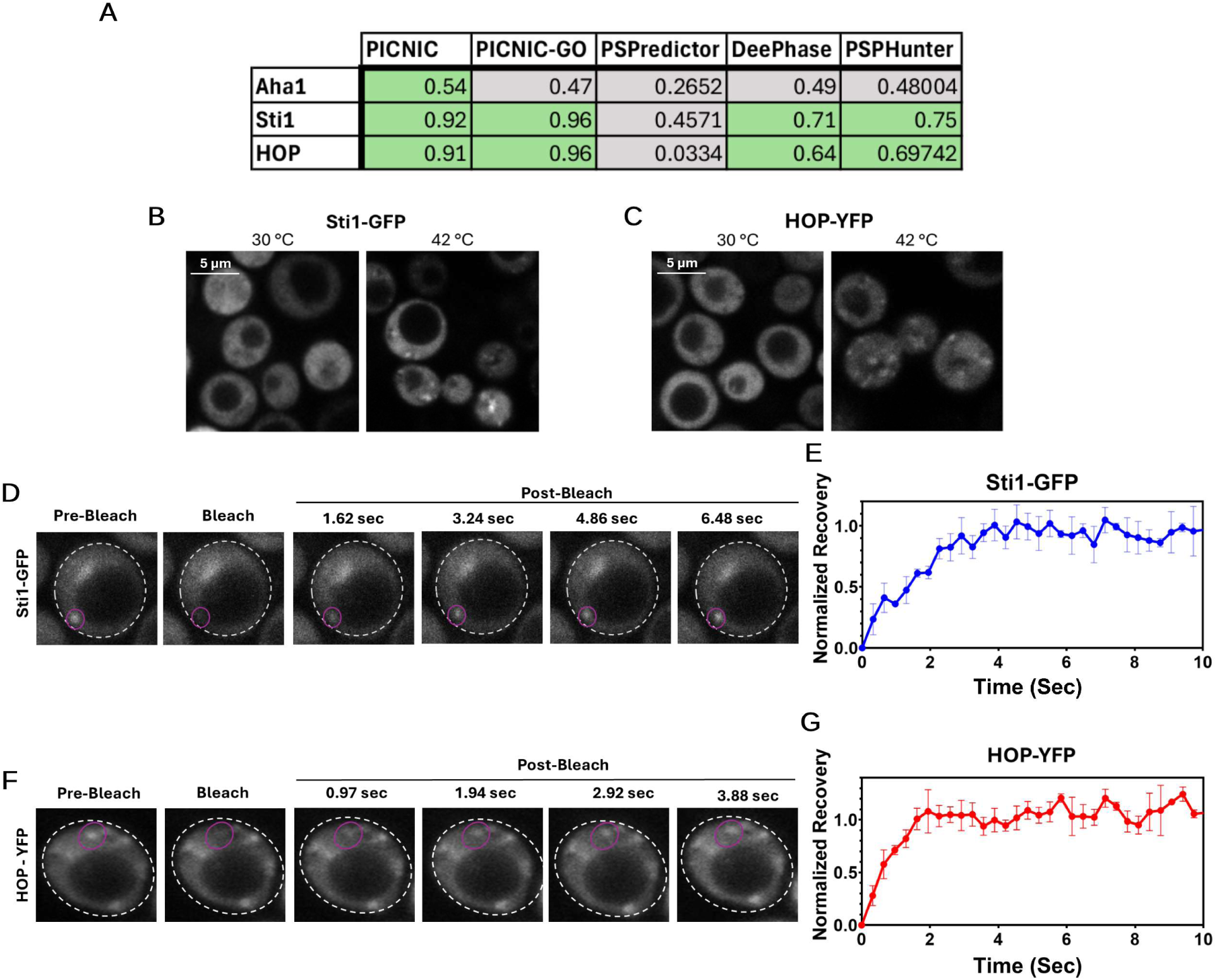
Sti1 and its human homologue HOP form dynamic condensates in the cytoplasm of yeast. (A) Condensation propensity of Sti1 (Uniprot ID: P15705), HOP (Uniprot ID: P31948), and AHA1 (Uniprot ID: Q12449) predicted by common LLPS prediction servers (scores above passing threshold shown in green). (B) Fluorescent microscopy of yeast expressing Sti1-GFP under its endogenous promoter before and after 42 °C heat shock. (C) Representative fluorescence microscopy images of *Δsti1* yeast cells expressing HOP-YFP before and after heat shock. (D) Representative FRAP images of a heat-shocked yeast cell expressing Sti1-GFP under its endogenous promoter before and after photobleaching. The bleach point is indicated by the purple line. (E) Quantification of the fluorescence recovery of photobleached Sti1-GFP foci. (F) Representative FRAP images of a heat-shocked *Δsti1* yeast cell expressing HOP-YFP before and after photobleaching. The bleach point is indicated by the purple line. (G) Quantification of the fluorescence recovery of photobleached HOP-YFP foci.

To determine whether the inclusions formed by Sti1 and HOP in response to protein misfolding stress exhibit liquid-like properties, we performed fluorescence recovery after photobleaching (FRAP) measurements. Heat shock was applied to induce the formation of Sti1-GFP foci in yeast cells (Figure 1B). HOP can functionally replace Sti1 in yeast[36], and we also observed that, identical to Sti1, heat shock induces the formation of cytoplasmic HOP-YFP foci when exogenously expressing HOP-YFP in *Δsti1* W303 yeast (Figure 1C). FRAP analysis of Sti1-GFP or HOP-YFP foci shows that the majority of cytoplasmic foci containing Sti1 or HOP are highly mobile and dynamic, exchanging rapidly with the surrounding environment (Figures 1D-G). These results indicate that stress-induced Sti1/HOP inclusions exhibit liquid-like properties, explaining their solubility and transient nature.

### HOP phase separates *in vitro*

We next sought to determine whether HOP actively drives protein condensation in cells or is simply recruited to other condensates containing misfolded proteins. To this end, we examined the phase separation properties of purified recombinant HOP *in vitro*. Since heat shock promotes the formation of HOP foci in cells, we monitored HOP across a range of temperatures. HOP solutions exhibited turbidity (OD_600_) at physiological temperatures (37 °C), with turbidity increasing as the temperatures rose. Brightfield microscopy reveals this increase in turbidity correlated with the appearance of droplets at higher temperatures (Figures 2A and B). Next, we assessed the effect of the molecular crowding agent PEG 3350, which facilitates phase separation of Hsp70 and Hsp90[25,37], on HOP. The addition of increasing concentrations of PEG causes spontaneous formation of HOP droplets at ambient temperature as observed by brightfield microscopy (Figure 2C) and fluorescent imaging using Alexa-680 conjugated to HOP (Figure 2D). During prolonged incubation, HOP droplets fuse upon contact, leading to an increase in average droplet size and a corresponding decrease in droplet number over time (Figures 2E-H), a well-established characteristic of liquid phase separation.

**Figure 2.**
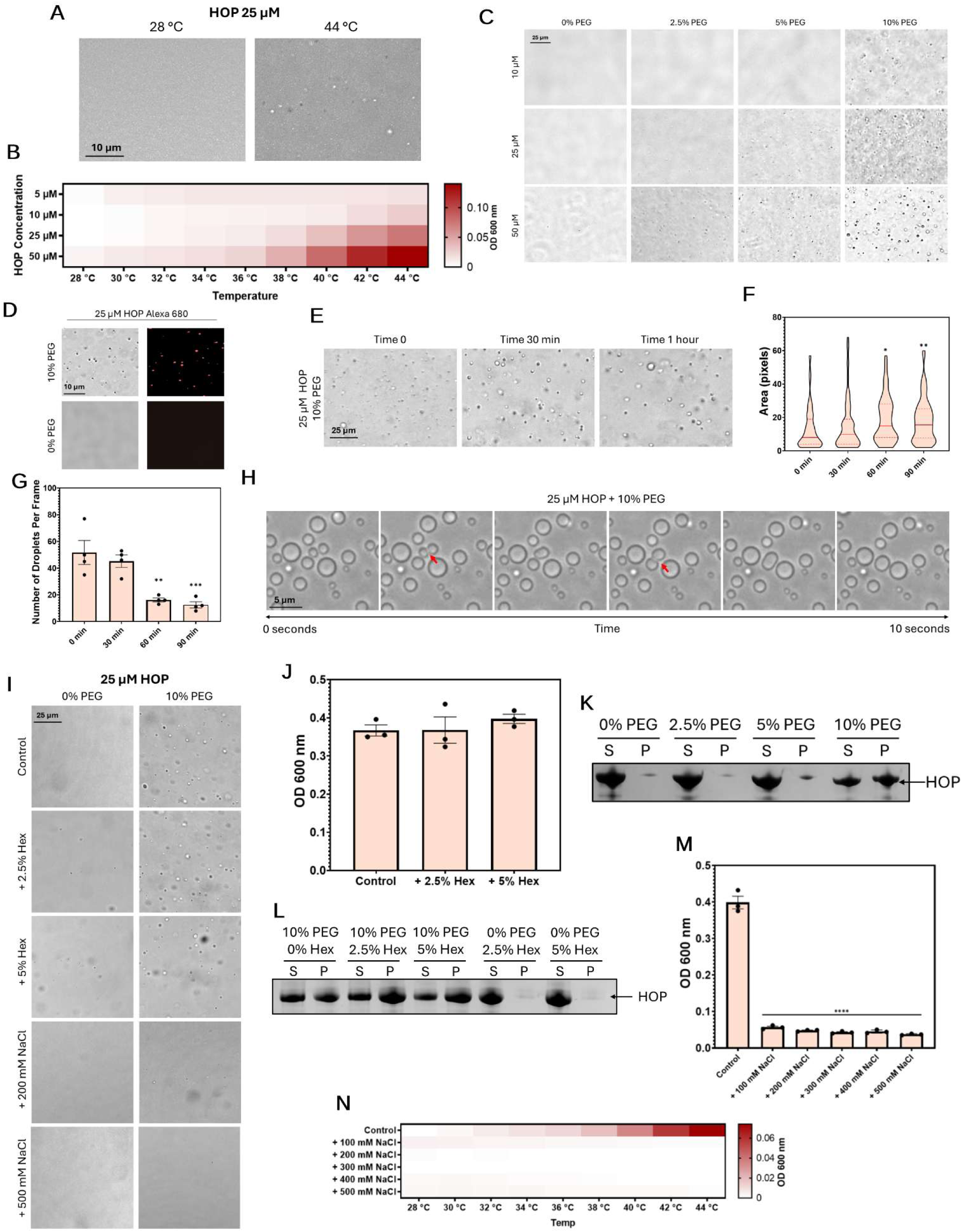
HOP forms condensates in vitro via electrostatic interactions. (A) Brightfield microscopy of 25 µM HOP condensates formed upon heating the solution from 28 °C to 44 °C. (B) Quantification of the solution turbidity at OD_600_ for a range of HOP concentrations over a series of temperatures. (C) Brightfield microscopy of various concentrations of HOP in a range of PEG concentrations at ambient temperature. (D) Brightfield and fluorescent microscopy of 25 μM Alexa 680 labelled HOP in 10% or 0% PEG. (E) Brightfield microscopy of 25 μM HOP in 10% PEG immediately after initiating condensation and after set time periods. (F-G) Quantification of the size (F) and number (G) of condensates observed for 25 μM HOP in 10% PEG at set time periods after initiating condensation. (H) Brightfield microscopy images showing an example of HOP condensate merging (point of fusion indicated by red arrow). (I) Brightfield microscopy of 25 μM HOP treated with various Hexanediol (Hex) and NaCl concentrations in the absence and presence of 10% PEG. (J) Quantification of the turbidity of 25 μM HOP treated with various Hexanediol concentrations in the presence of 10% PEG. (K) Sedimentation profile of 25 μM HOP in the presence of various PEG concentrations. The aqueous phase/supernatant is indicated by “S” and the condensate phase/pellet is indicated by “P”. (L) Sedimentation profile of 25 μM HOP treated with various Hexanediol concentrations in the absence and presence of 10% PEG. (M) Quantification of the turbidity of 25 μM HOP treated with various NaCl concentrations in the presence of 10% PEG. (N) Quantification of the turbidity of 25 μM HOP treated with various NaCl concentrations over a range of temperatures. To determine statistical significance, unpaired one-way ANOVA tests were used to compare means and standard deviations between relevant controls and experimental data sets (each data set was composed of a minimum of three replicas). Statistical significance is represented by an asterisk, where **** is P < 0.0001, *** is P < 0.001, ** is P < 0.01, and * is P < 0.05. Error bars represent standard errors of the mean.

### Electrostatic interactions drive HOP phase separation

Next, we examined the molecular mechanisms underlying HOP phase separation. Both hydrophobic[38,39] and electrostatic[19,27] interactions are commonly implicated in driving the phase separation of polypeptides[40]. The aliphatic alcohol 1,6-hexanediol (Hex) disrupts phase separation driven by hydrophobic groups but does not affect phase separation mediated by electrostatic interactions[41]. Treatment with both 2.5% and 5% Hex, concentrations sufficient for disrupting condensates mediated by hydrophobic interactions[41], had no significant effect on the turbidity or droplets observable by microscopy in HOP solutions (Figures 2I and J). To further verify that Hex did not affect the proportion of HOP in protein droplets, HOP solutions were subject to centrifugation, and the condensate phase (pellet) was separated from the aqueous phase (supernatant). The presence of 10% PEG caused a significant amount of HOP to partition into the condensate phase, and the addition of Hex failed to reduce this proportion (Figures 2K and L). Taken together, these results suggest that the HOP droplets observed *in vitro* are not primarily driven by hydrophobic interactions.

To determine if the stress-dependent Sti1-GFP foci observed *in vivo* share similar Hex-insensitivity of HOP droplets formed *in vitro*, yeast cells were treated with Hex and Sti1-GFP foci were monitored. It has been previously demonstrated that treatment of yeast with Hex dissolves P granules and Fus condensates, but the more solid-like stress granules are not susceptible to Hex treatment[42]. We subjected yeast cells expressing Sti1-GFP to heat shock and then incubated at optimal growth temperatures in media containing various concentrations of Hex. Interestingly, treatment with 2.5% and 5% Hex not only failed to dissolve Sti1 foci but instead resulted in a moderate increase in their abundance during the recovery period (Supplemental Figure 1). The results suggest that Sti1-GFP foci are not driven by hydrophobic interactions. Instead, Hex may promote Sti1-GFP phase separations either through direct effects or as a consequence of the cytotoxicity associated with prolonged exposure[42]. The Hex insensitivity of Sti1 and HOP droplets we observe both *in vitro* and *in vivo* indicates that the HOP condensation is not driven by hydrophobic interactions.

Hex-insensitive condensates can be formed by charge peptides that are highly sensitive to the addition of salt, e.g., NaCl[41]. To investigate whether electrostatic interactions drive the phase separation of HOP, we examined the effect of increasing NaCl concentrations. The addition of salt significantly reduced both the turbidity and number of droplets in HOP solutions containing 10% PEG (Figures 2I and M). To assess whether the temperature-induced phase separation of HOP exhibits similar sensitivity to NaCl, we monitored the turbidity of HOP solutions containing increasing NaCl concentrations across increasing temperatures. Increasing concentrations of NaCl suppressed the increase in turbidity at temperatures above 37 °C (Figure 2N). The insensitivity of HOP phase separation to Hex and strong sensitivity to NaCl suggests that HOP phase separation is mainly driven by electrostatic interactions.

### HOP condensation depends on multiple domain interactions

Previous *in vitro* studies on engineered proteins containing tandem TPR repeats demonstrate how TPR domains can promote condensate formation[19]. Given the presence of multiple TPR domains in HOP, we speculated that they are key to HOP phase separation. To predict which regions of HOP drive condensation, we employed PSPHunter, a machine-learning prediction tool that identifies amino acid residues critical for protein phase separation[35].

PSPHunter predicts three regions of driving residues, one in the TPR1 domain (residues 47-67), one adjacent to the TPR2A domain (residues 189-221), and one within the TPR2A domain (residues 286-308) (Figure 3A), which aligns well with our hypothesis that the TPR domains may drive phase separation of HOP.

**Figure 3.**
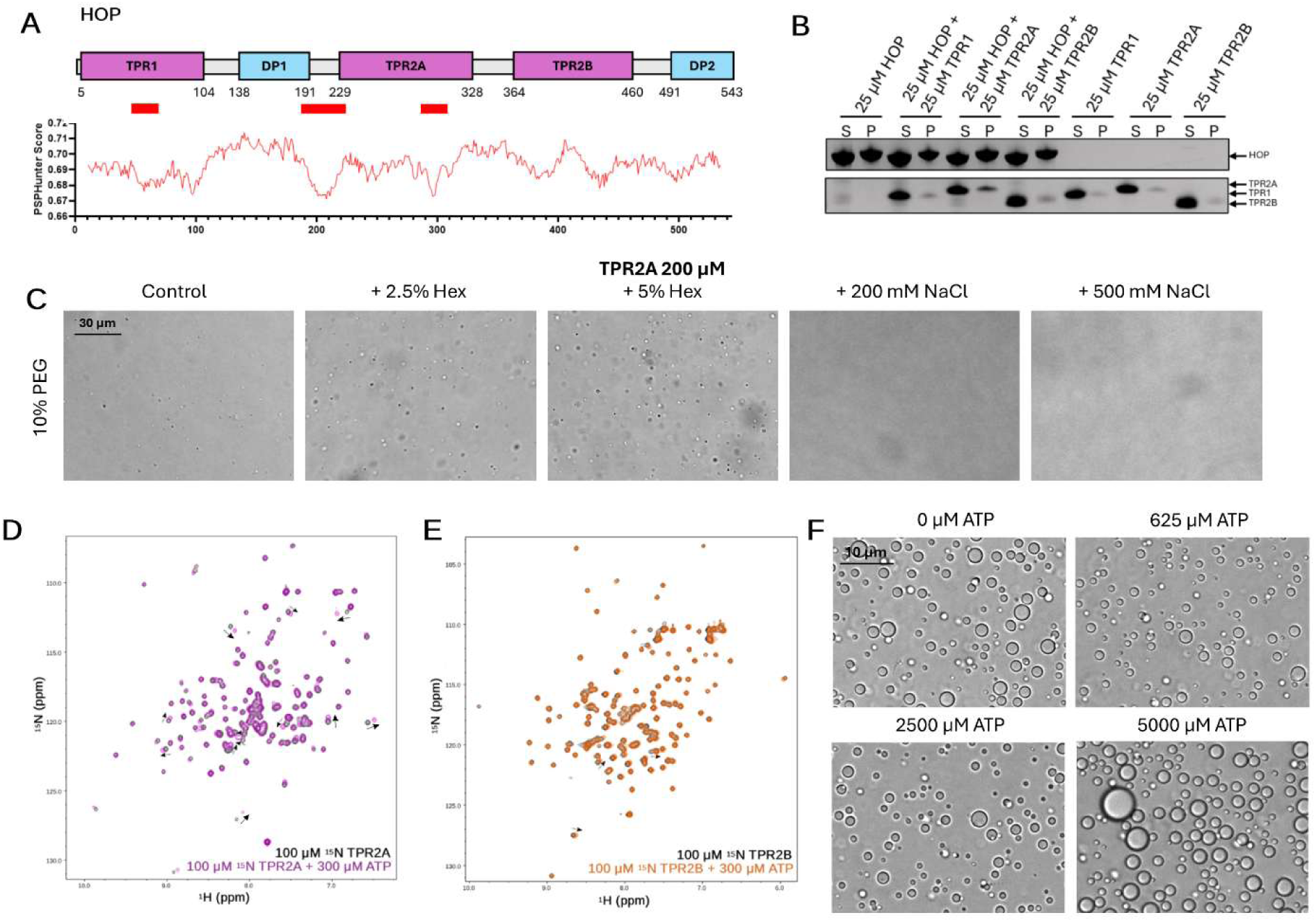
Intermolecular interactions with the TPR2A domain drive HOP condensation. (A) Domain map of HOP with predicted driver residues indicated in Red and corresponding PSPHunter score depicted below. (B) Sedimentation profile of 25 μM TPR1, TPR2A, and TPR2B in the absence and presence of equimolar HOP in 10% PEG. The aqueous phase/supernatant is indicated by “S” and the condensate phase/pellet is indicated by “P”. (C) Representative brightfield microscopy images of 200 μM TPR2A in the absence and presence of various concentrations of Hexanediol (Hex) and NaCl in 10% PEG. (D) NMR analysis of ^15^N-labeled TPR2A at 100 µM in the absence (black) and the presence of 300 µM ATP (Purple) with largest observable CSPs indicated by arrows. (E) NMR analysis of ^15^N-labeled TPR2B at 100 µM in the absence (black) and the presence of 300 µM ATP (Orange) with largest observable CSPs indicated by arrows. (F) Representative brightfield microscopy images of 25 μM HOP in the absence or presence of various concentrations of ATP in 10% PEG.

Longer segments of TPR repeats show greater propensities for phase separation than individual TPRs[19,27]. We thus sought to determine whether all three TPRs of HOP or any individual TPR domains drive its phase separation. Solutions of TPR1, TPR2A, or TPR2B in the presence of 10% PEG were monitored for droplet formation. Markedly higher protein concentrations (>300 μM for TPR1, >200 μM for TPR2A and TPR2B) were required to observe droplets of the TPR domains in comparison to full-length HOP, which condensates at as low as 10 μM. Furthermore, the droplets formed by the individual TPR domains are smaller in size than those formed by full-length HOP (Supplemental Figure 2). The substantial reduction in phase separation observed for individual TPR domains suggests that HOP condensation involves interactions across multiple TPR domains.

Given that no single TPR domain recapitulated the efficient phase separation propensity of full-length HOP, we hypothesized that transient TPR interactions collectively contribute to its droplet formation. NMR ^1^H-^15^N HSQC experiments were used to screen for interactions between the individual TPR domains and full-length HOP. No significant spectral changes were observed upon the addition of equimolar HOP to either ^15^N labelled TPR1 or TPR2B (Supplemental Figure 3), indicating no strong interactions between the separate TPR domains and full-length HOP. By contrast, the addition of equimolar HOP to ^15^N labelled TPR2A resulted in the broadening of many peaks, plausibly reflecting transient interactions between TPR2A and HOP that contribute to phase separation (Supplemental Figure 4).

To determine whether the interaction between separate TPR domains and full-length HOP is prevalent during phase separation, solutions of the TPR domains in the absence and presence of equimolar full-length HOP and with the addition of PEG were subjected to centrifugation to isolate the condensate phase. In the absence of full-length HOP, the TPR domains were predominantly present in the soluble phase. The addition of HOP had no notable effect on the partitioning of TPR1 or TPR2B but increased the proportion of TPR2A in the condensate phase (Figure 3B), indicating that the transient interaction between TPR2A and full-length HOP recruits TPR2A into HOP droplets. Similar to full-length HOP, increasing concentrations of NaCl reduce the TPR2A droplet formation, suggesting that electrostatic interactions mediate TPR2A phase separation (Figure 3C). The addition of increasing concentrations of Hex increases the observable TPR2A droplets, further suggesting that hydrophobic interactions do not drive TPR2A phase separation (Figure 3C). The increased phase separation of TPR2A upon the addition of Hex could be due to Hex facilitating ionic-dependent condensation, as has been observed for the Hex-facilitated condensation of chromatin *in vitro*[43]. The similarities between TPR2A and full-length HOP phase separation suggest a predominant role of TPR2A in driving HOP condensation.

### ATP promotes HOP phase separation

HOP can directly interact with the adenine group of ATP via the TPR1-TPR2A segment of HOP (residues 1-359)[44]. The condensate phase of positively charged protein sequences complements the ability of ATP to bridge π-interactions as an amphiphilic hydrotrope, thereby enhancing phase separation[45–47]. We speculated that ATP may enhance HOP phase separation by bridging intermolecular interactions between its charged TPR domains. To identify potential ATP binding interfaces, we performed NMR ^1^H-^15^N HSQC experiments to screen for interactions between the individual ^15^N-labelled TPR domains and ATP. Only minor spectral changes were observed for ^15^N-labelled TPR1 upon the addition of 3-fold molar excess of ATP (Supplemental Figure 5). By contrast, more notable peak shifts were observed in several hydrophobic residues of TPR2A and to a lesser extent TPR2B (Figures 3D and E). These observations suggest that TPR1 and TPR2B transiently interact with ATP, whereas TPR2A binds more strongly. Notably, some of the residues showing the chemical shift changes in TPR2A upon the addition of ATP are adjacent to the carboxylic clamp residues, which are crucial for Hsp90 binding (Supplemental Figure 6). The interaction between the TPR2A domain and the hydrophobic adenine group of ATP could mask hydrophobic regions on the TPR2A while bridging electrostatic interactions via the negatively charged hydrophilic triphosphate moiety, similar to the ATP-driven phase separation of Nonannotated P-body dissociating polypeptide[46].

To test whether the interactions between ATP and the TPR2A and TPR2B domains of HOP enhance HOP phase separation, we monitored solutions containing HOP and increasing ATP concentrations. HOP phase separation increased with increasing ATP concentrations, with the addition of 5 mM ATP producing large HOP droplets (Figure 3F). Taken together, these results suggest that the presence of ATP enhances HOP phase separation, potentially through its interactions with the TPR2A and TPR2B domains.

### Y354E HOP exhibits reduced phase separation in vitro

The elongated shape frequently adopted by proteins containing tandem TPR repeats allows folded domains to engage in multivalent interactions along the TPR scaffold[19]. The tandem TPR2A and TPR2B domains in HOP also adopt an elongated structure due to the highly structured linker connecting these two domains[20]. We speculated that post-translational modifications affecting the conformation of the TPR2A-TPR2B domains modulate HOP phase separation. To test this, we utilized the Y354E amino acid substitution to mimic the charge change caused by phosphorylation of Y354 within the linker between TPR2A and TPR2B[48]. This phosphomimetic destabilizes the structured linker, causing TPR2A-TPR2B to adopt more dynamic conformations, in which the two TPR domains are disconnected[36].

We observe that, upon addition of 10% PEG, solutions of Y354E HOP spontaneously become turbid and form droplets. Time course measurements of turbidity following 10% PEG addition reveal that Y354E HOP exhibits significantly lower turbidity than wild type HOP. Notably, unlike wild type HOP, which reaches a peak turbidity that remains stable, the turbidity of Y354E HOP reaches a lower peak around 1 hour after PEG addition and then progressively decreases after 2 hours (Figure 4A). Isolating the condensate phase of aged HOP and Y354E HOP solutions by centrifugation reveals that the proportion of both wild type and Y354E HOP in the condensate phase increases over time. However, there is a marked decrease in the proportion of Y354E in the condensate phase after 20 minutes and 2 hours that is not observed for wild-type HOP (Figure 4B). Taken together, these results suggest that the Y354E amino acid substitution has a reduced propensity to phase separate compared to wild type HOP.

**Figure 4.**
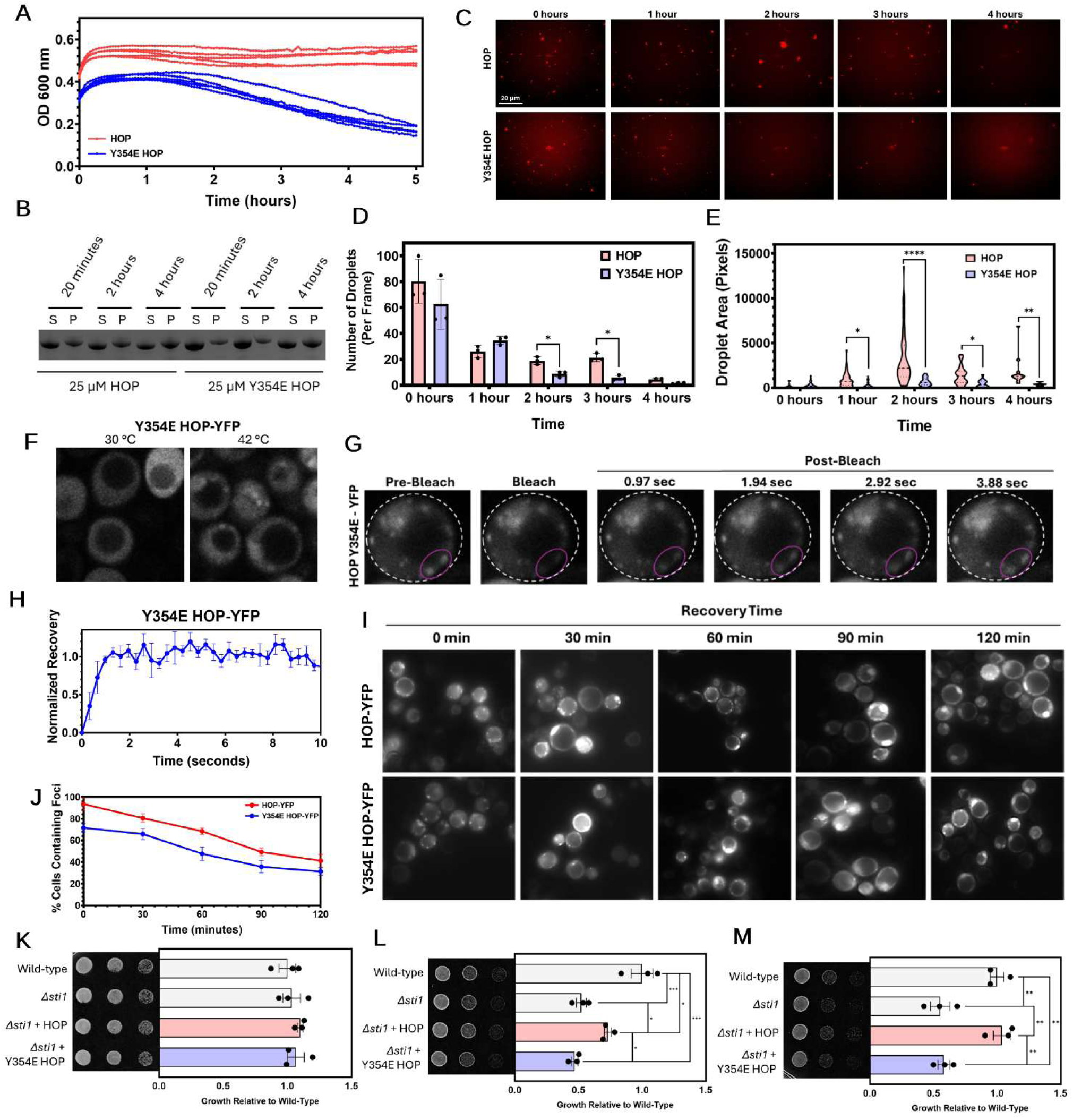
The Y354E substitution reduces HOP condensation and impairs the cell’s ability to recover from heat shock. (A) Quantification of turbidity for HOP and Y354E HOP in 10% PEG immediately after initiating condensation and for a prolonged period. (B) Sedimentation profile of HOP and Y354E HOP in 10% PEG after set time periods. The aqueous phase/supernatant is indicated by “S” and the condensate phase/pellet is indicated by “P”. (C) fluorescent microscopy images of Alexa 680 labelled HOP or Y354E HOP in 10% PEG immediately after initiating condensation and after set time periods. (D-E) Quantification of the number (D) and size (E) of HOP and Y354E HOP condensates immediately after initiating condensation and after set time periods. (F) Representative fluorescence microscopy images of *Δsti1* yeast cells expressing Y354E HOP-YFP before and after heat shock. (G) Representative FRAP images of a heat-shocked *Δsti1* yeast cell expressing Y354E HOP-YFP before and after photobleaching. The bleach point is indicated by the purple line. (H) Quantification of the fluorescence recovery of photobleached Y354E HOP-YFP foci. (I) Fluorescent microscopy images of *Δsti1* yeast cells expressing either HOP-YFP or Y354E HOP-YFP immediately after heat shock and during recovery. (J) Quantification of the percent of cells containing either HOP-YFP or Y354E HOP-YFP foci during recovery. (K-M) Growth assay of control yeast and of *Δsti1* yeast cells expressing HOP or Y354E HOP grown at 30 °C (K), 37 °C (L), or heat shocked for 1 hour at 42 °C before being grown at 30 °C (M). To determine statistical significance, unpaired one-way or two-way ANOVA tests were used to compare means and standard deviations between relevant controls and experimental data sets (each data set was composed of a minimum of three replicas). Statistical significance is represented by an asterisk, where **** is P < 0.0001, *** is P < 0.001, ** is P < 0.01, and * is P < 0.05. Error bars represent standard errors of the mean.

Next, we investigated the size and abundance of Y354E HOP droplets and compared them to wild-type HOP. To increase the accuracy of size measurements, HOP and Y354E HOP were labelled with an Alexa 680 dye and droplets were monitored by fluorescent microscopy. We observe no significant difference between wild type and Y354E HOP in droplet number or size immediately after initiating phase separation (Figure 4C). Both HOP and Y354E droplets increase in size and decrease in number over time (Figures 4D-E), suggesting that both wild type HOP and Y354E droplets effectively fuse upon contact. However, at all subsequent time points after initiating phase separation, the average size of the Y354E HOP droplets was significantly smaller than that of HOP. Furthermore, we observe a significant reduction in the number of Y354E HOP droplets in comparison to HOP at the 2- and 3-hour time points (Figures 4D and E). In agreement with the turbidity measurements, the reduced droplet size and number of Y354E HOP suggest that this phosphomimetic substitution reduces the propensity for HOP phase separation.

### The Y354E substitution in HOP causes fewer cytoplasmic liquid-like foci in cells during stress

We next expressed Y354E HOP-YFP in *Δsti1* W303 yeast cells to determine whether similar effects can be observed *in vivo.* Expression of Y354E HOP-YFP in *Δsti1* W303 yeast produces similar stress-dependent foci as seen with HOP-YFP (Figure 4F; see Figure 1C for comparison). FRAP measurements of these foci show that they rapidly exchange with the surrounding environment, similar to wild-type HOP-YFP (Figures 4G and H; see Figures 1F and G for comparison). However, compared to wild type HOP-YFP, significantly fewer Y354E HOP-YFP expressing cells contained cytoplasmic foci immediately following heat shock and during the recovery (Figures 4I and J). Taken together, these results suggest that Y354E exhibits a reduced propensity to form condensates within cells compared to wild type HOP.

### The Y354E Phosphomimetic substitution impairs stress-related functions of HOP

Since the Y354E phosphomimetic substitution reduced the phase separation propensity of HOP both *in vitro* and in yeast cells, we sought to determine whether Y354E impairs HOP functions under protein misfolding stress. To this end, we assessed the growth of wild-type and *Δsti1* W303 yeast expressing wild type HOP or Y354E HOP. The deletion of STI1 causes no growth defect at 30 °C but impairs yeast growth under protein misfolding stress[3,49].

Consistent with these findings, we observed no significant differences in growth among all tested yeast strains under optimal temperature conditions (30°C, Figure 4K). However, when the yeast cells were grown under mild heat stress (37 °C), simulating chronic misfolding stress, the expression of wild-type HOP in *Δsti1* W303 yeast partially complemented its growth defect associated with the deletion of STI1, whereas Y354E HOP did not (Figure 4L). To align this growth defect with reduced phase separation, we heat shocked yeast cells (42 °C for one hour) before spotting them on plates that were incubated at 30 °C. After the heat shock, the expression of wild-type HOP in *Δsti1* W303 yeast completely complemented the growth defects caused by the deletion of STI1, while *Δsti1* W303 yeast expressing Y354E HOP showed no significant growth improvement compared to controls lacking STI1 (Figure 4M). Due to the diminished phase separation observed for Y354E HOP both in cells and *in vitro*, we deduce that the disruption of phase separation contributes to impaired function of Y354E HOP during misfolding stress. Taken together, these results suggest that the Y354E phosphomimetic substitution impairs the ability of HOP to perform phase separation, and thus has a diminished capacity to protect cells from stress.

### Hsp70 and Hsp90 regulate the formation and the dissolution of Sti1 condensates

Sti1/HOP appears to be central to the formation of foci that sequester misfolded proteins during stress and coordinate cellular proteostasis, as the recruitment of Hsp90, Hsp70, and Hsp104 to proteasome-dependent heat-induced inclusions requires Sti1[6]. In addition, the clearance of Sti1-GFP foci after the alleviation of stress is, at least partially, dependent on the presence of other cellular chaperones[3]. We previously observed that inhibition of Hsp82 (yeast homologue of Hsp90) significantly impaired dissolution of Sti1-GFP foci during the recovery from heat shock[3]. To determine whether the levels of Hsp82 and Ssa1 (a yeast homologue of Hsp70) alter the dissolution of Sti1-GFP foci, we overexpressed either Hsp82 or Ssa1 and measured the number of Sti1-GFP foci following heat shock. Neither Hsp82 nor Ssa1 overexpression induced Sti1-GFP condensation in the absence of heat stress (Supplemental Figure 7). Fewer cells overexpressing Hsp82 contained Sti1-GFP condensates immediately following heat shock and during the recovery period compared to wild-type cells. Conversely, the overexpression of Ssa1 impaired the dissolution of Sti-GFP condensates as compared to wild-type cells (Figures 5A and B). Taken together, these results suggest that both Hsp70 and Hsp90 regulate the formation and dissolution of Sti1-GFP condensates in cells.

**Figure 5.**
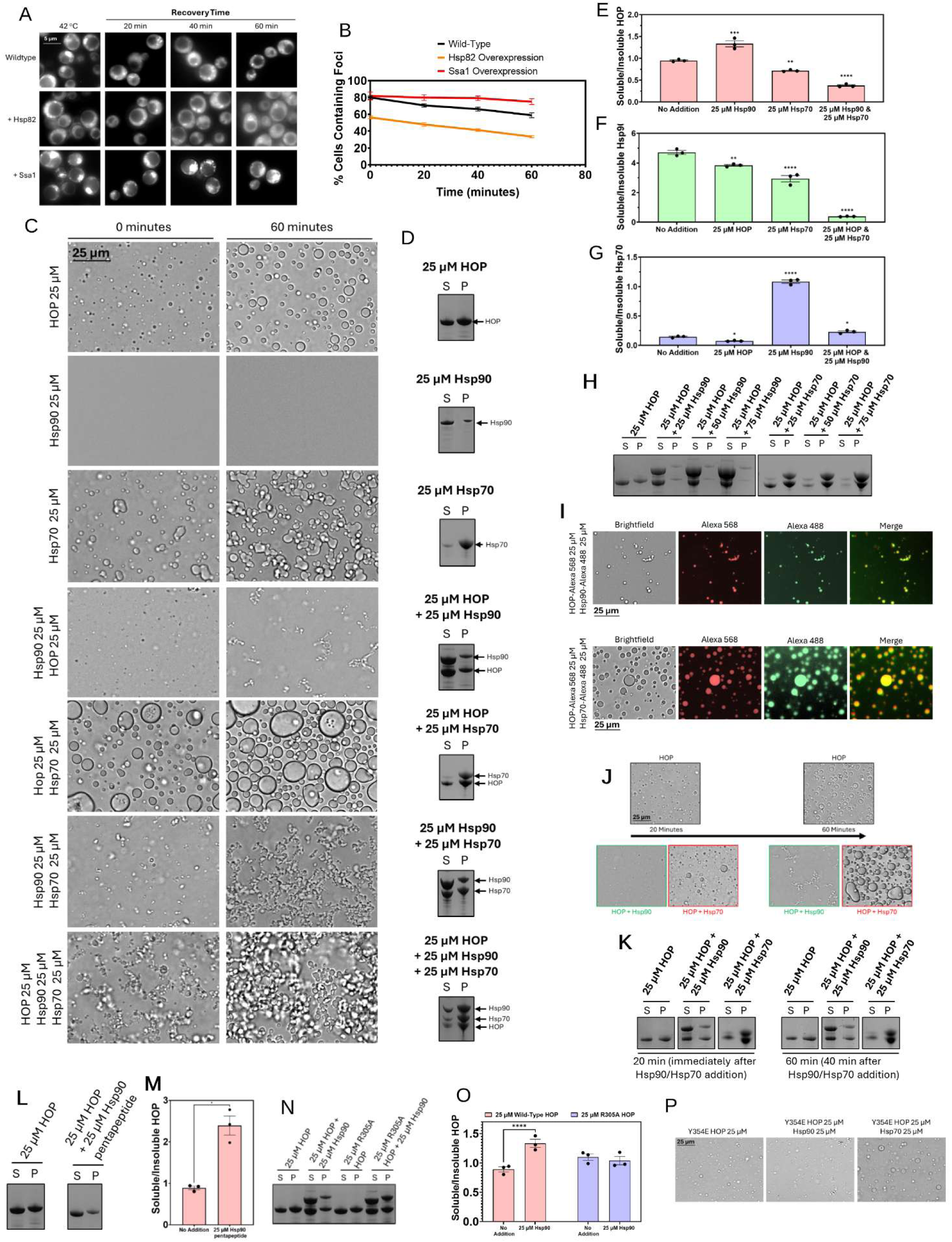
Hsp90 and Hsp70 have opposing effects on HOP phase separation. (A) Representative fluorescence microscopy images of yeast cells expressing Sti1-GFP under its endogenous promoter and transformed with overexpression plasmids of either Hsp82 or Ssa1 after heat shock and during recovery. (B) Quantification of the percent of yeast cells expressing Sti1-GFP under its endogenous promoter and transformed with overexpression plasmids of either Hsp82 or Ssa1 that contain Sti1 foci after heat shock and during recovery. (C) Representative brightfield microscopy images of 25 μM HOP, Hsp90, and Hsp70 in various combinations in the presence of 10% PEG. (D) Sedimentation profile of 25 μM HOP, Hsp90, and Hsp70 in various combinations in the presence of 10% PEG. The aqueous phase/supernatant is indicated by “S” and the condensate phase/pellet is indicated by “P”. (E-G) Quantification of the HOP (E), Hsp90 (F), and Hsp70 (G) sedimentation profiles in various combinations of HOP, Hsp90, and Hsp70 in 10% PEG. (H) Sedimentation profile of 25 μM HOP with increasing concentrations of Hsp90 or Hsp70 in 10% PEG. (I) Representative brightfield and fluorescent microscopy of 25 μM HOP labelled with Alexa-568 in the presence of equimolar Hsp90 labelled with Alexa-488 or Hsp70 labelled with Alexa-488. (J-K) Representative brightfield microscopy images (J) and sedimentation profile (K) of solutions containing 25 μM HOP in the absence or presence of equimolar Hsp90 or Hsp70 and 10% PEG. Solutions of HOP were incubated for 20 minutes at room temperature and then either equimolar Hsp90 or Hsp70 was added to the solutions containing existing HOP droplets. The solutions of 25 μM HOP in the absence or presence of either equimolar Hsp90 or Hsp70 were then incubated for an additional 40 minutes. (L) Sedimentation profile of 25 μM HOP in the absence or presence of equimolar Hsp90 pentapeptide in 10% PEG. (M) Quantification of the sedimentation profile of 25 μM HOP in the absence or presence of equimolar Hsp90 pentapeptide in 10% PEG. (N) Sedimentation profile of 25 μM HOP and 25 μM R305A HOP in the absence or presence of equimolar Hsp90 in 10% PEG. (O) Quantification of the sedimentation profile of 25 μM HOP and 25 μM R305A HOP in the absence or presence of equimolar Hsp90 in 10% PEG. (P) Brightfield microscopy images of 25 μM Y354E HOP in the presence of equimolar Hsp90 or Hsp70 in 10% PEG. To determine statistical significance, unpaired one-way or two-way ANOVA tests were used to compare means and standard deviations between relevant controls and experimental data sets (each data set was composed of a minimum of three replicas). Statistical significance is represented by an asterisk, where **** is P < 0.0001, *** is P < 0.001, ** is P < 0.01, and * is P < 0.05. Error bars represent standard errors of the mean.

### HOP, Hsp90, and Hsp70 have different phase separation propensities

Since the relative levels of Hsp82 and Ssa1 had significant effects on the Sti1 condensation observed *in vivo*, we sought to determine how Hsp90 and Hsp70 modulate HOP phase separation *in vitro*. Both Hsp90[37] and Hsp70[25] have previously been shown to undergo phase separation on their own *in vitro.* Accordingly, Hsp70 formed condensates under the same conditions we used for our experiments with purified HOP; however, no Hsp90 phase separation was observed (Figure 5C). Notably, Hsp90 was previously shown to phase separate only with 20% PEG 3350[37], suggesting that it may require more severe molecular crowding than HOP and Hsp70 to condensate. Centrifugation revealed that the majority of Hsp70, but only a small proportion of Hsp90, was present in the condensate phase (Figure 5D), which may likely be due to minor amount of Hsp90 unfolding and precipitation during incubation rather than phase separation.

### Hsp90 reduces HOP phase separation in vitro

To further examine how Hsp90 affects HOP phase separation, solutions of HOP were monitored in the presence of equimolar amounts of Hsp90. The presence of Hsp90 reduced the amount of visible HOP droplets and impaired their fusion upon contact, evidenced by the persistence of dispersed clusters of small droplets after 60 minutes of incubation (Figure 5C). Hsp90 also significantly decreased HOP partitioning into condensates (Figures 5C, D, and E), in a concentration-dependent manner (Figure 5H). Reduced HOP phase separation in the presence of Hsp90 suggests that Hsp90 facilitates the dissolution of HOP droplets. To determine whether Hsp90 is associated with HOP within condensates, HOP solutions labelled with an Alexa568 dye were mixed with Hsp90 solutions labelled with an Alexa488 dye. The fluorescent signals of HOP and Hsp90 overlap within the observed droplets, confirming their colocalization (Figure 5I). Furthermore, the proportion of Hsp90 in the condensate phase significantly increases in the presence of HOP, as compared to Hsp90 alone (Figures 5D and F), suggesting that a fraction of Hsp90 is associated with HOP droplets.

To determine whether Hsp90 removes HOP from condensates, we initiated HOP phase separation, and Hsp90 was added after 20 minutes. The addition of Hsp90 immediately decreased both the size of HOP condensates and the proportion of HOP in the condensate phase. After an additional 40 minutes, Hsp90 further reduced the size of the observed condensates (Figure 5J) but did not substantially decrease the amount of HOP in condensates as compared to after the initial addition of Hsp90 (Figure 5K). Taken together, these results suggest that Hsp90 associates with existing HOP condensates and removes HOP from the condensate phase.

### Hsp90 dissolves HOP droplets through its interaction with the TPR2A domain of HOP

We next examined whether the Hsp90-binding site within the TPR2A domain is required for removing HOP from condensates. To test this, we used the minimal Hsp90 pentapeptide (MEEVD), which binds the carboxylate clamp of TPR2A (K229, N233, N264, K301, and R305)[50]. Addition of the Hsp90 pentapeptide to solutions of HOP significantly reduced the proportion of HOP in the condensate phase (Figures 5L and M). Increasing concentrations of the pentapeptide further reduced the size of condensate droplets and promoted clustering of droplets that fail to merge, similar to the effects of full-length Hsp90 (Supplemental Figure 8). Notably, the addition of the Hsp90 pentapeptide to solutions of the TPR2A domain also diminished the number of protein droplets (Supplemental Figure 8). These findings suggest that occupying the Hsp90-binding site on the TPR2A domain disrupts critical interactions or induces conformational changes in HOP resulting in reduced HOP phase separation.

The R305A amino acid substitution in the TPR2A domain of HOP reduces its Hsp90 binding affinity[51]. We used R305A to investigate whether efficient TPR2A binding is required for Hsp90-mediated dissociation of HOP droplets. No significant difference in HOP condensation was observed upon introduction of the R305A substitution (Supplemental Figure 8). Furthermore, adding the Hsp90 pentapeptide to solutions of R305A HOP failed to reduce droplet number, size, or fusion (Supplemental Figure 8). These results confirm that the Hsp90 pentapeptide disrupts HOP condensation through direct interactions with the TPR2A domain.

Similarly, the addition of equimolar Hsp90 significantly reduced the proportion of wild-type HOP in the condensate phase but had no effect on R305A (Figures 5N and O). Taken together, these results suggest that the direct interaction between Hsp90 and the TPR2A domain of HOP is required for Hsp90-mediated removal of HOP from condensates.

### Hsp70 enhances HOP phase separation in vitro

Next, we tested the effects of Hsp70 on the phase separation of HOP. In contrast to Hsp90, the addition of equimolar Hsp70 produces larger condensate droplets than those observed for either Hsp70 or HOP alone. The droplets of Hsp70 and HOP together continue to fuse upon contact, producing much larger droplets after 60 minutes of incubation than each protein alone (Figure 5C). Furthermore, Hsp70 significantly increases HOP partitioning into the condensate phase (Figures 5D and E) in a concentration-dependent manner (Figure 5H). Interestingly, in the presence of HOP, a larger fraction of Hsp70 was also observed to partition into the condensate phase (Figures 5D and G). The increased proportion of both HOP and Hsp70 within condensates (Figures 5D, E, and G) indicates that the larger condensates contain greater amounts of both proteins, rather than reflecting a mere change in droplet morphology. Mixing Alexa568-labelled HOP with Alexa488-labelled Hsp70 reveals overlapping fluorescent signals within the observed condensate droplets, confirming that both proteins are present within the same large droplets (Figure 5I). The synergistic phase separation of HOP and Hsp70 suggests that each protein promotes the condensation of the other.

To determine whether Hsp70 can also enlarge preformed HOP condensates, Hsp70 was added to solutions of HOP 20 minutes after the initiation of phase separation. The addition of Hsp70 increased the size of HOP condensate droplets and the proportion of HOP in the condensate phase. After an additional 40 minutes of incubation, Hsp70 continued to increase the size of the droplets (Figure 5J), although the proportion of HOP in the condensate phase only increased slightly as compared to immediately upon Hsp70 addition (Figure 5K). Taken together, these results suggest that Hsp70 can associate with existing HOP condensates and increase their size.

### Interplay between HOP, Hsp90 and Hsp70 in phase separation

Since Hsp90 and Hsp70 exert opposing effects on HOP condensation, we next sought to investigate the effect of both chaperones combined on HOP condensation. To this end, we first examined the effect of Hsp90 on Hsp70 condensation. The presence of Hsp90 reduced the Hsp70 droplets and suppressed their fusion upon contact (Figure 5C). Centrifugation-based analysis of the condensate phase revealed a substantially higher proportion of Hsp90 and a lower proportion Hsp70 (Figures 5D, F, and G) in condensates compared to condensates formed by Hsp90 or Hsp70 alone.

We next monitored the condensation of solution mixtures combining HOP, Hsp90, and Hsp70. Many droplets were formed, which progressively fused into larger droplets over the 60 minutes incubation (Figure 5C). Notably, isolation of the condensate phase revealed a significantly larger proportion of HOP in the presence of equimolar Hsp90 and Hsp70 than in the addition of only Hsp70 (Figures 5C and D). Furthermore, these condensates contained significantly more Hsp90 than any other combination (Figures 5D and F) but substantially less Hsp70 than those observed for Hsp70 alone or for equimolar HOP-Hsp70 mixtures (Figures 5D and G). The shift of HOP and Hsp90 toward the condensate phase indicates that Hsp70 enhances their propensity to condensate. Moreover, the changes in droplet appearance compared to HOP-Hsp70 mixtures suggests that the trimolecular combination may form liquid droplets with a distinct morphology.

To further evaluate how relative concentrations of Hsp90 and Hsp70 regulate HOP condensation, solution mixtures of HOP, Hsp90, and Hsp70 were monitored in three-fold higher concentration of either Hsp90 or Hsp70 to HOP. Excess of Hsp90 significantly decreased the proportion of HOP, Hsp90, and Hsp70 in the isolated condensates (Figures 6A and B). In contrast, excess of Hsp70 substantially increased the fraction of all three proteins partitioning into the condensate phase (Figures 6A and B). These opposing effects of Hsp90 and Hsp70 suggest that the phase separation of the HOP-Hsp90-Hsp70 mixture exists in a delicate equilibrium, where the relative abundance of each component modulates droplet formation.

**Figure 6.**
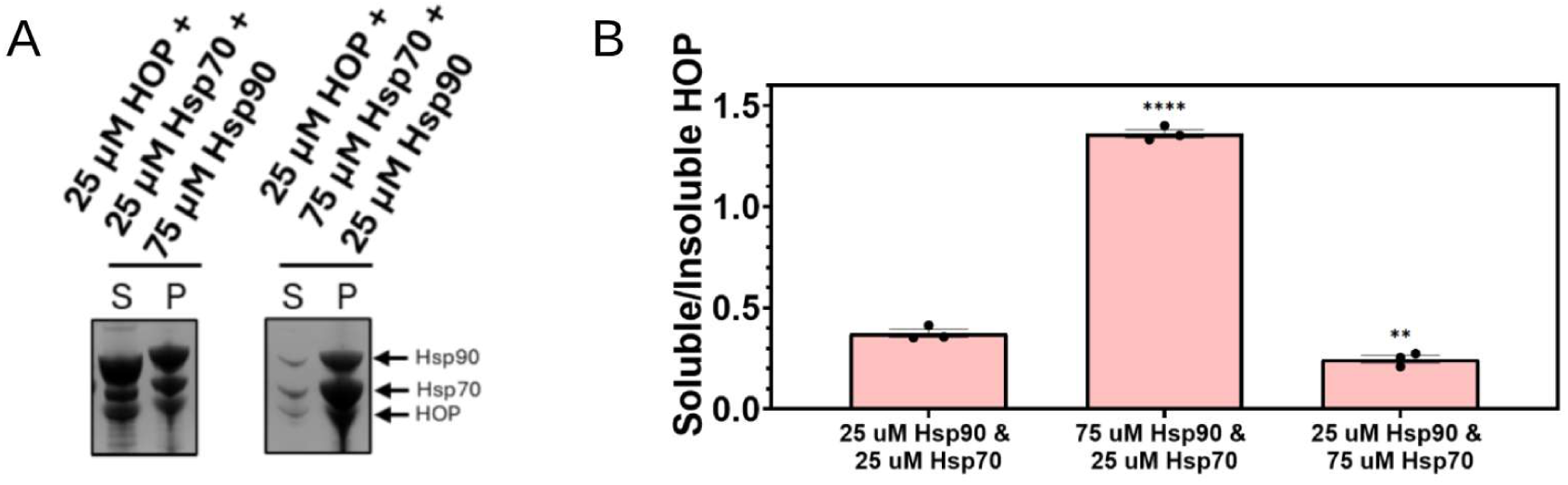
The relative ratios of each member of the HOP-Hsp70-Hsp90 mixture impacts HOP phase separation. (A-B) Sedimentation profile (A) and quantification (B) of 25 μM HOP with equimolar Hsp70 and excess Hsp90 or equimolar Hsp90 and excess Hsp70 in 10% PEG. The aqueous phase/supernatant is indicated by “S” and the condensate phase/pellet is indicated by “P”. To determine statistical significance, an unpaired one-way ANOVA tests were used to compare means and standard deviations between relevant controls and experimental data sets (each data set was composed of a minimum of three replicas). Statistical significance is represented by an asterisk, where **** is P < 0.0001, *** is P < 0.001, ** is P < 0.01, and * is P < 0.05. Error bars represent standard errors of the mean.

### ATP promotes phase separation of HOP and Hsp90

Since the binding of ATP alters the conformations and activities of both Hsp90 and Hsp70[1,52,53], we investigated if ATP changes Hsp90 and Hsp70-mediated HOP condensation. The addition of increasing ATP concentrations caused no changes to the Hsp70 droplets; however, the presence of 5 mM ATP enhanced Hsp90 phase separation (Supplemental Figure 9, Figures 7A-C). The addition of 5 mM ATP to solutions of equimolar HOP and Hsp90 increased the number of droplets and the proportion of HOP in the isolated condensate phase (Figures 7A, B, and D). Notably, the addition of 5 mM ATP to solutions of equimolar HOP and Hsp70 had no effect on the phase separation, nor on the proportion of HOP and Hsp70 in the condensate phase (Figures 7A, B, and D). Taken together, these results suggest that 5 mM ATP is sufficient to enhance the phase separation of HOP and Hsp90 and suppresses the Hsp90-mediated removal of HOP from condensates. Conversely, we observe no significant effect of ATP addition on Hsp70 phase separation, and the addition of Hsp70 abolishes the effect of ATP on HOP phase separation.

**Figure 7.**
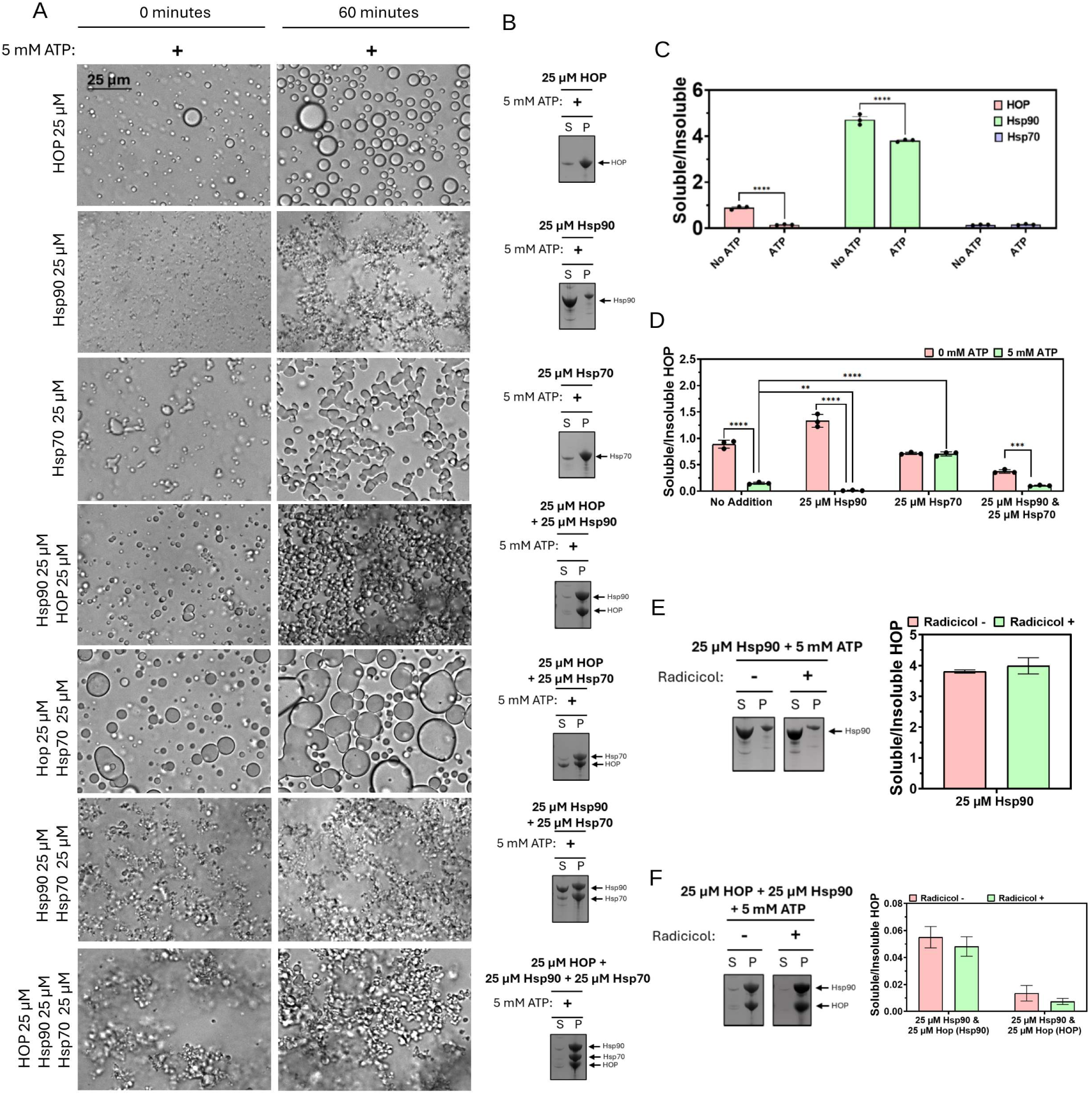
The addition of ATP promotes phase separation of HOP independent of hydrolysis. (A) Representative brightfield microscopy images of 25 μM HOP, Hsp90, and Hsc70 in various combinations in the presence of 5 mM ATP and 10% PEG. (B) Sedimentation profile of 25 μM HOP, Hsp90, and Hsp70 in various combinations in the presence of 5 mM ATP and 10% PEG. The aqueous phase/supernatant is indicated by “S” and the condensate phase/pellet is indicated by “P”. (C) Quantification of the sedimentation profiles of HOP, Hsp90, or Hsp70 in the absence (data from Figure 5 for statistical comparison) and presence of 5 mM ATP in 10% PEG. (D) Quantification of the HOP sedimentation profile in the absence and presence of Hsp90, Hsp70, or both Hsp90 and Hsp70, in the absence (data from Figure 5 for statistical comparison) and presence of 5 mM ATP in 10% PEG. (E) Sedimentation profile and quantification of 25 μM Hsp90 with the addition of 5 mM ATP and in the absence or presence of Radicicol. (F) Sedimentation profile and quantification of 25 μM HOP and equimolar Hsp90 with the addition of 5 mM ATP, and in the absence or presence of Radicicol. To determine statistical significance, unpaired one-way or two-way ANOVA tests were used to compare means and standard deviations between relevant controls and experimental data sets (each data set was composed of a minimum of three replicas). Statistical significance is represented by an asterisk, where **** is P < 0.0001, *** is P < 0.001, ** is P < 0.01, and * is P < 0.05. Error bars represent standard errors of the mean.

Next, we investigated if ATP alters the condensation of combined HOP-Hsp90-Hsp70. The addition of 5 mM ATP to solutions of equimolar Hsp90 and Hsp70 increased droplet formation and prevented Hsp90 from reducing the proportion of Hsp70 in the condensate phase (Figures 7A and B). The addition of 5 mM ATP to HOP, Hsp90, and Hsp70 combined significantly increased the proportion of HOP, Hsp90 and Hsp70 in the condensate phase (Figures 7A, B, and D). Overall, we observe that the addition of 5 mM ATP increases HOP and Hsp90 phase separation and inhibits Hsp90-mediated reduction of HOP and Hsp70 condensation, suggesting that the cellular levels of ATP may alter the effectiveness of Hsp90-mediated regulation of HOP condensation.

### Hydrolysis is not required for ATP mediated Hsp90 phase separation

We next sought to determine whether ATP facilitates Hsp90 phase separation via ATP hydrolysis by Hsp90 or by merely acting as a hydrotrope that bridges intermolecular interactions, a mechanism previously observed for other proteins[46,47]. Radicicol, which occupies the ATP binding pocket of Hsp90 and thereby reduces its ATPase activity[54], was added in two-fold excess to solutions of Hsp90 and 5 mM ATP. The addition of Radicicol did not significantly reduce the proportion of Hsp90 in the condensate phase (Figure 7E), suggesting that the Hsp90 droplets formed in the presence of 5 mM ATP does not require ATPase activity. Furthermore, when added to solutions of Hsp90, HOP, and 5 mM ATP, the excess of Radicicol had no significant effect on the proportion of HOP in the isolated condensate phase (Figure 7F). The lack of Radicicol sensitivity suggests that the Hsp90 ATPase activity does not reduce the proportion of HOP in the condensate phase in the presence of 5 mM ATP. This indicates that the ATP hydrotrope activity accounts for the effects on HOP-Hsp90-Hsp70 phase separation, rather than Hsp90-mediated ATP hydrolysis.

## Discussion

We report that the stress-dependent cytoplasmic foci of Sti1/HOP are liquid-like, resembling other stress-induced cellular compartments formed by liquid-liquid phase separation. Our biophysical analysis demonstrates that HOP readily undergoes phase separation, forming protein droplets resembling the foci observed in cells. Relative concentrations of Hsp90 and Hsp70 regulate the extent of HOP phase separation both *in vitro* and *in vivo*, with Hsp90 reducing HOP phase separation and Hsp70 driving HOP droplet formation. Our results suggest that the interaction between Sti/HOP, Hsp90, and Hsp70 regulate the incorporation of these molecular chaperones into condensate compartments during misfolding stress.

Phase separation has predominantly been proposed to be driven by intrinsically disordered regions (IDRs), due to their ability to form weak multivalent interactions. However, it is now recognized that the multivalency of multi-domain proteins comprised of tandem structured repeats linked by IDRs makes for ideal candidates for phase separation[15,19]. Specifically, the phase separation of proteins comprised of tandem TPR domain repeats, such as Rapsn[27], is driven by electrostatic interactions involving multiple TPR repeats[19,27]. Similarly, we observe that the phase separation of HOP is disrupted by increasing salt concentrations, suggesting that electrostatic interactions drive HOP phase separation. Furthermore, the individual TPR domains of HOP display reduced phase separation propensity, suggesting that the phase separation of HOP cannot be attributed to any single domain, but rather is driven by electrostatic interactions spanning multiple TPR domains. The TPR2A domain was the only TPR domain for which transient interactions were observed with full-length HOP. These transient interactions were sufficient to draw TPR2A into HOP condensates, suggesting TPR2A may play a notable role in driving the condensation of HOP.

The elongated conformation adopted by proteins containing tandem repeat TPR domains is key for facilitating their phase separation[19]. Structural analysis has demonstrated that the structured linker connecting the tandem TPR2A and TPR2B domains causes HOP to adopt an elongated structure[20]. We observe that destabilizing this structured linker with the Y354E amino acid substitution[36], mimicking phosphorylation of Y354, significantly reduces the phase separation of HOP both *in vitro* and *in vivo*. In particular, the Y354E substitution significantly reduces the HOP-YFP condensates observed following heat shock in comparison to wild-type HOP. Furthermore, while expression of wild-type HOP in *Δsti1* yeast completely complements for the growth reduction observed following heat shock, expression of Y354E HOP fails to complement the deletion of STI1. The formation of stress induced Sti1 foci is crucial for the coordinating the Hsp90, Hsp70, Hsp104 and the proteasome[6,55], and facilitating the clearance of soluble misfolded proteins[3]. The inability of Y354E HOP to complement the deletion of STI1 in yeast following heat shock suggests that the stress related functions of HOP may be at least to some extent reliant on the phase separation propensity of HOP.

Sti1/HOP bridges Hsp90 and Hsp70, facilitating the complex formation and the refolding cascade[1,56]. However, Hsp90 and Hsp70 can also engage with multiple cochaperones[57–59] or form binary complexes through direct interactions[55,60], suggesting redundancy across cochaperones in their ability to form Hsp90/Hsp70 complexes. Of note, recent research suggests that cochaperones play larger roles in coordinating complex proteostasis networks[61]. Sti1/HOP has a more direct role during proteostasis stress by sequestering soluble misfolded proteins into cytoplasmic foci in a Hsp90 independent manner[3]. Sti1/HOP protein levels are tightly regulated, with overexpression inducing the formation of Sti1 foci even in the absence of stress stimuli[3]. Notably, we observe that altering the relative expression of the major chaperone partners of HOP, Hsp90 and Hsp70, regulate Sti1 foci formation. Consistent with our previous studies showing that inhibition of Hsp90 activity in yeast impairs the clearance of Sti1 foci during recovery from cellular stress[3], we find that overexpression of Hsp90 significantly reduces the number of stress-induced Sti1 foci. In contrast, overexpression of Hsp70 in yeast prevents the proper clearance of Sti1 foci following stress. Both Hsp90 and Hsp70 also regulate the phase separation of HOP *in vitro*, with Hsp90 dissolving HOP droplets and Hsp70 co-condensing with HOP into larger droplets. It is important to note that the regulatory effects of Hsp70 may be bidirectional, as HOP also influences Hsp70 condensation to favor formation of larger droplets These regulatory effects are also observed when Hsp90 and Hsp70 are added to existing HOP droplets, indicating that HOP retains its interactions with chaperone partners in the condensates. The regulatory effects of Hsp90 can be attributed to the direct interaction with TPR2A, as the minimal Hsp90 pentapeptide (MEEVD), which binds TPR2A, is sufficient to disrupt the phase separation of both HOP and the TPR2A domain alone. Furthermore, the R305A substitution in HOP, previously shown to reduce Hsp90 binding, negates the influence of Hsp90 addition on HOP phase separation, highlighting the potential importance of TPR2A domain for HOP phase separation.

Hsp70 has previously been observed forming large oligomeric assemblies when in the presence of substrates[62,63]. The addition of HOP restructures these assemblies, allowing Hsp70 to transfer substrates to Hsp90[63]. This aligns with the significant morphological changes observed when Hsp70 condensates form in the presence of HOP. It has been speculated that these previously observed Hsp70 assemblies exist due to high chaperone to substrate ratios[63], however, we propose that these assemblies may also be driven by the phase separation propensities of Hsp70 and HOP. This phase separation propensity allows HOP to form condensates in rapid response to acute stress, in contrast to other quality control condensates, such as JUNQ and IPOD, which require the accumulation of misfolded proteins and not molecular chaperones[64]. HOP can interact with misfolded substrates[23], and the stress induced Sti1 foci co-localize with misfolded protein markers in yeast cells[3], suggesting that HOP may recruit misfolded substrates into the rapidly forming condensates.

It is also relevant to note that in mammalian cells HOP is a critical regulator of abnormal assemblies of chaperones and clients, known as the epichaperome[65], which is triggered by cell stress in cancer[65] and protein misfolding in neurodegenerative diseases[66]. We anticipate that the ability to form phase separation and its regulation by Hsp70 and Hsp90 are also critical for epichaperome formation.

A potential emerging picture is that during acute stress, HOP and Hsp70 co-condensate, acting as a first line of defense to recruit misfolded substrates and sequestering them for effective processing. Specifically, this immediate response by HOP and Hsp70 may facilitate the triage of misfolded proteins to subsequent proteostasis pathways, potentially explaining the accumulation of soluble ubiquitinated proteins observed upon the deletion of Sti1/HOP[3]. It has recently been demonstrated that nutrient deficiency induces the formation of cytoplasmic Sti1 foci resembling those observed upon heat stress[67]. Chen et al. propose that during nutrient deprivation, the intrinsic phase separation capacity of Sti1 causes it to form condensates in conjunction with Hsp70 and other proteostasis chaperones[67]. This aligns with our hypothesis that HOP and Hsp70 co-condensation primes the proteins for effective clearance by Hsp90 once proteostatic stress subsides, thus determining the transient nature of these condensates.

Together, our research reveals the molecular mechanisms by which phase separation facilitates HOP to coordinate the proteostasis response during stress and how the relative abundance of HOP, Hsp70, Hsp90, and likely other members involved in the proteostasis network, appears to regulate the formation and clearance of the stress induced HOP condensates. This previously understudied feature of HOP provides new insights into how HOP sequesters misfolded proteins during acute stress. Furthermore, the phase separation propensity of HOP deserves consideration in the complex relationship between HOP and protein misfolding diseases. HOP interacts with numerous disease-associated proteins, including TAR DNA-binding protein 43 (TDP-43)[68], prion protein[4], amyloid-β[69], and α-synuclein[23], and promote the formation of cytotoxic α-synuclein oligomers[70]. Soluble aggregates of α-synuclein, and other disease-associated proteins, are normally cleared via the proteosome system[71], suggesting that they may be sequestered by HOP within stress-dependent condensates. Of note, increased levels of HOP can also affect and worsen the misfolding of amyloid-β[69] and TDP-43[72]. Therefore, interactions within these condensates may represent a maladaptive mechanism through which HOP contributes to the accumulation of toxic oligomers in disease contexts.

## Declarations

### Availability of data and material

All study data are included in the article and/or Supplemental Information.

## Acknowledgments

Research reported in this publication was supported by funding from Schulich School of Medicine and Dentistry, Canada Foundation for Innovation (CFI), Ontario Research Fund (ORF), and NSERC Research Tools and Instruments Awards to Western University through use of the BioCORE Facility.

## Funding

B.S.R. is supported by a Canada Graduate Research Scholarship, M.L.D. is supported by a Discovery Grant (RGPIN 2024-05867) from the Natural Sciences and Engineering Research Council (NSERC), W.Y.C is supported by a Discovery Grant (RGPIN-2025-05599) from NSERC. M.A.M.P. received support from the Canadian Institutes of Health Research (CIHR, PJT 162431, PJT 195987), NSERC (RGPIN 2021-03592), BrainsCAN Canada First Research Excellence Fund Accelerator Awards (Initiative for Translational Neuroscience), Canada Foundation for Innovation, Ontario Research Fund, Alzheimer’s Association (USA), Weston Brain Institute, Weston Family Foundation, as well as support from New Frontiers Research Fund ( NFRFT-2022-00051, TRIDENT). M.A.M.P. is a Tier I Canada Research Chair in Neurochemistry of Dementia.

## Authors’ contributions

B.S.R., designed and performed research, analyzed data, generated figures, and wrote the paper. B.E.G. and M.V. performed research and analyzed data. S.H., D.P.A and P.L. contributed reagents. M.A.M.P, W.Y.C, and M.L.D designed research, contributed reagents or analytical tools, and wrote the paper. All authors aided in revising the manuscript’s intellectual content. All authors read and approved the submitted manuscript.

## Competing interests

The authors declare no competing interests.

